# Prebiotic Potential of Non-digestible Oligosaccharides Isolated from Barnyard Millet (*Echinochloa frumentacea*) Grain

**DOI:** 10.64898/2026.02.26.708187

**Authors:** Sachin Maji, Paramita Biswas, Shivangi Agrawal, Sandip Shit, Satyahari Dey

## Abstract

It is encouraging to see the growing acceptance of millet among the general public, driven by its numerous health benefits. Various research efforts have focused on the many non-digestible oligosaccharides (NDOs) in millets, given their significant nutraceutical potential. Barnyard millet is a viable candidate for extracting NDOs owing to its superior nutritional value and affordability. In the present study, crude oligosaccharides were extracted from barnyard millet under optimized conditions: 80% aqueous ethanol (v/v) at a 1:5 sample-to-solvent ratio for 1 h at 60 °C, yielding 3.61±0.24%. Extracted oligosaccharides were further purified through adsorption chromatography and dialyzed through dialysis membranes (500 Da). Several significant peaks at m/z values of 527, 689, 852, 1014, and 1338 were detected in the MALDI-TOF mass spectrum, indicating that barnyard millet oligosaccharides (BMOs) comprise various oligosaccharides with degrees of polymerization (DP) from 3 to 8. BMOs were treated with digestive enzymes (porcine pancreatic and salivary α-amylase) and an artificial acidic solution, and results showed that approximately 91% of BMOs were not digested. The positive prebiotic scores and the generation of lactic and short-chain fatty acids (SCFAs) upon fermentation by various probiotic strains indicate the prebiotic potential of BMOs. In this study, the results also showed that the presence of BMOs increased auto-aggregation of lactobacilli and enhanced the adhesion of probiotics to HCT116 cells. Our findings indicate that, with its nutritional benefits, barnyard millet may serve as a viable reservoir of beneficial carbohydrates, including NDOs.

**Highlights:**

- Extraction of BMOs under optimized parameters.
- Partial purification of BMOs through charcoal column chromatography and dialysis.
- A positive prebiotic activity score conferred the prebiotic potential of BMOs.
- Production of short-chain fatty acids by fermentation of BMOs.
- BMOs increased the auto-aggregation percentage of different probiotics.

## 1. Introduction

The meaning and concept of food are changing with the development of science and technology, from feeding people’s survival needs to keeping them healthy and disease-free. Nutraceuticals and functional foods are among the trending food concepts in the past few decades. NDOs have entered the nutraceutical league due to their health benefits and prebiotic effects. NDOs are typically considered prebiotics because they have a lower degree of polymerization (DP 3–10) and are more readily soluble. These two factors together make them better at changing the composition of gut microbes. Prebiotics are dietary nutrients that are not digested or broken down by digestive enzymes. They also do not get absorbed in the upper gastrointestinal system, so they reach the colon intact. Within the colon, they selectively promote, in particular, the proliferation and metabolic activity of specific helpful microbial populations, such as *Lactobacillus* and *Bifidobacterium* species, ultimately contributing to the enhancement of host gastrointestinal health and overall well-being (Macfarlane et al., 2006).

A result of oligosaccharide digestion induced by anaerobic gut microbiota is the production of short-chain fatty acids (SCFAs) (Asadpoor et al.,2021), primarily propionate, butyrate, and acetate (Cummings et al., 1981, 1995). Short-chain fatty acids enhance gastrointestinal health by maintaining a strong protective layer of the intestinal wall, stimulating mucus secretion (Silva et al., 2020), and providing various health benefits, such as anti-inflammatory, anticancer, antidiabetic, and neuroprotective properties (Xiong et al., 2022). The production of SCFAs makes the colon acidic, which helps good probiotic bacteria flourish and prevents harmful pathogens from growing (Mondal et al., 2022). The auto-aggregation assay is an important *in vitro* screening tool used to evaluate the potential of probiotic bacteria to colonize the GI tract (Cozzolino et al., 2020). When oligosaccharides (such as FOS, GOS, or XOS) are present, probiotic cell-surface properties are selectively altered, often making it easier for them to form multicellular aggregates that facilitate their adherence and retention in the host ecosystem (Yin et al., 2024). In the presence of prebiotics, an increasing percentage of auto-aggregation of probiotics indicates greater colonization, the primary step in forming a mature biofilm.

Cereal grains fulfill the dietary fibre, protein, energy, minerals, and vitamins requirements for human health, which is the maximum percentage of the world’s food production. Hence, it becomes an abundant and economically viable source of prebiotics owing to its content of specific non-digestible carbohydrate molecules. Millets are unique among cereal grains, as they have a better nutritional profile, greater health benefits, and remarkable environmental sustainability than major staple foods like rice and wheat. Barnyard millet (*Echinochloa sp*.) is one of the fast-growing minor millets widely cultivated in Asia’s temperate regions (Renganathan et al., 2020; Nithiyanantham et al., 2019). According to reports, from 2015 to 2018, India became the largest producer of barnyard millet (IIMR, 2018). Barnyard millet has a wide variety, and among them, the two most common are *Echinochloa frumentacea* (Indian variety) and *Echinochloa esculenta* (Japanese variety) (Sood et al., 2015). Currently, experts are showing interest in barnyard millet due to its health benefits, including antidiabetic (Ugare et al., 2011), anticancer (Ramadoss & Sivalingam, 2019), antioxidant (Watanabe et al., 1999), and immunostimulatory (Srinivasan et al., 2019) properties. It is a rich source of carbohydrates, protein, fiber, and micronutrients such as iron (Fe) and calcium (Singh et al., 2009). According to Manju et al. (2025), the world dietary fibre market value could reach USD 19.97 billion by 2034. India, as a major producer of millets, colored rice, barley, etc., offers great promise for developing prebiotics from these sources that are affordable for underprivileged populations. Generally, the non-digestible carbohydrates extracted from millets, which have prebiotic potential, are either polysaccharides or enzyme-degraded oligosaccharides (Palaniappan et al., 2017; Devi et al., 2011). Mondal et al. (2022) reported the direct extraction and purification of oligosaccharides from pearl millet and their prebiotic efficacy. The specific oligosaccharides extracted from barnyard millet have not yet been documented, however. In the current study, an effort is made to extract and purify the oligosaccharides from barnyard millet and explore their prebiotic efficacy.

## 2. Materials and methods

### 2.1 Materials and microorganisms

Barnyard millet (*Echinochloa frumentacea*) (product name: nilgiris barnyard kuthiravali, article no. 40002439, manufactured and packed by Future Consumer Limited) was purchased from Big Bazar (Kharagpur, India). 99.99% (v/v) ethanol (made in China) was procured from a local vendor; de-ionized water was taken from a Milli-Q (German) system; all other solvents and 98% (v/v) sulfuric acid were from Merck (German). 3,5-dinitrosalicylic acid (SRL), potassium sodium tartarate (Merck), phenol (Fisher Scientific), activated charcoal powder (Loba Chemie Pvt. Ltd.), celite 545 (Merck, Germany), silica gel 60 F_254_ plates (Merck), Orcinol (SRL), DMEM (Gibco), FBS (Thermo Fisher Scientific), trypsin-EDTA (Himedia), cell culture antibiotics (Thermo Fisher Scientific) were purchased from a local supplier. All carbohydrate standards (glucose, sucrose, maltose, raffinose, FOS), and pancreatic α-amylase, salivary α-amylase were from Sigma–Aldrich Co. Ltd. General laboratory salts, culture media components, including MRS broth, were obtained from Himedia (India). Dialysis tube (MWCO 500 Da), and all short-chain fatty acid standards were imported from Spectrum Laboratories (USA) and Dr. Ehrenstorfer (Germany), respectively, through a local supplier. All other chemicals used were analytical grade.

The probiotic strains *Lactobacillus rhamnosus* MTCC 1408, *Lactobacillus acidophilus* MTCC 10307, and *Lactobacillus plantarum* MTCC 2621 were ordered from MTCC, India, and the enteric control strain *Escherichia coli* ATCC 15223 was used in all experiments. The human colorectal cancer cell line (HCT 116) was ordered from the National Centre for Cell Sciences (NCCS), Pune, India.

### 2.2 Extraction of BMOs

The millet grains were rinsed twice with water and then dried at 50 °C for 12 h. After cleaning, the grains were crushed in a grinder, and the ground flour was sifted through different meshes (size between 1000 and 125 μm). Most of the particles fell between 500 and 150 μm in size. The extraction procedure of barnyard millet oligosaccharides was optimized (not discussed in this article), and the maximum yield was obtained by treating grain flour with 80% aqueous ethanol (v/v) with a 1:5 sample solvent ratio for 1 h at 60 °C. After extraction, the slurry was spun at 4 °C with 10,000 g for 30 mins, and the supernatant was collected. The ethanol was removed from the collected extract using a rotary evaporator. The concentrated sample was used for further experiments.

### 2.3 Quantification of sugar and identification of carbohydrates via thin-layer chromatography (TLC)

The total sugar and reducing sugar concentrations in the sample were measured using the phenol-sulfuric acid assay (Dubois et al., 1956) and the DNSA method (Miller, 1959), respectively. TLC was performed on silica 60 F_254_ plates as the stationary phase to identify the number of repeating monomer units of the sugars in the crude extract. Acetic acid, butanol, and water in the ratio 1:2:1 were used as the mobile phase, and orcinol-sulfuric acid was used as the spraying reagent.

### 2.4 Partial purification of extracted oligosaccharides

Adsorption column chromatography was performed to purify the crude oligosaccharide extract. Using varying percentages (v/v) of ethanol (0 - 90%) as the mobile phase, crude extracted oligosaccharides were partially purified using an activated charcoal-celite (1:1) column (Nobre et al., 2011). Ethanol was separated out from eluted fractions using a rotary evaporator after the chromatography process, and the required fractions were then dialysed using a dialysis membrane (MWCO 500 Da). To visualize the sugar profiles of each fraction and the dialyzed sample, a TLC was performed using the method mentioned in 2.3. The presence of phenolic compounds was checked by TLC using a silica 60 F_254_ plate as the stationary phase, and a mobile phase prepared at a 6:5:1 ratio of formic acid, ethyl acetate, and toluene was used for chromatographic separation (Lawag et al., 2023). Following the separation, the plates were seen under UV light at 254 nm (Stanek & Jasicka-Misiak, 2018) and by the unaided eye after staining with a 2% (w/v) ethanolic ferric chloride solution (Jović et al., 2023). The presence of phenolic compounds was validated by comparing the retention factors (R_f_ values) of the separated bands with those of known phenolic standards. Protein concentration was quantified after purification using the Bradford Method (Hammond & Kruger, 2003).

### 2.5 Profiling of BMOs by MALDI-TOF

To chemically characterise a pure oligosaccharide, its molecular weight must be precisely determined to understand the molecule’s DP. After extraction, the molecular mass of each oligosaccharide was identified by MALDI-TOF MS (Bruker Ultraflextreme MALDI-TOF/TOF). It was performed using the method described in Liu and Rochfort (2015) with slight modifications. Before dialysis, 1 mg of the extracted lyophilized sample was dissolved in 0.1 mL of 0.01 N NaCl solution. 20 μL of 2,5-dihydroxybenzoic acid (DHB) matrix was added to 20 μL of the oligosaccharide-containing NaCl solution. 2 μL of that mixed solution was placed on a polished steel anchor plate (Bruker Daltonics) and permitted to dry prior to MALDI-TOF analysis. Spectra were acquired within the m/z range of 200 to 2000.

### 2.6 Thermal properties of BMOs

To understand the thermal stability and physicochemical properties of BMOs, Thermogravimetric Analysis/Differential Thermal Analysis (TGA/DTA) was carried out. Thermal analysis was performed using a Thermogravimetric/Differential Thermal Analyser (PerkinElmer, USA). In an alumina pan within a nitrogen environment, the sample (BMOs) was heated from 35 to 600 °C at a rate of 10 °C/min.

### 2.7 Prebiotic potentiality of BMOs

#### 2.7.1 Non-digestibility assay of BMOs

Non-digestibility indicates food that is not digested in the human digestive tract. An *in vitro* digestion assay was carried out by adding the sample to each phosphate-buffered solution containing salivary α-amylase and porcine pancreatic α-amylase, respectively. Acid resistance of BMOs was tested by treating the samples with 160 mM HCl. Each reaction mixture containing 1%(w/v) sample (BMOs) was incubated at 37±1 °C for 6 h. To assess the total and reducing sugar content, a 1 mL sample from each reaction tube was collected at the starting time and after 6 h of hydrolysis, followed by enzyme deactivation (Mondal et al., 2022). FOS and potato starch were used as controls. The following equation was used to figure out the percentage breakdown of the samples:

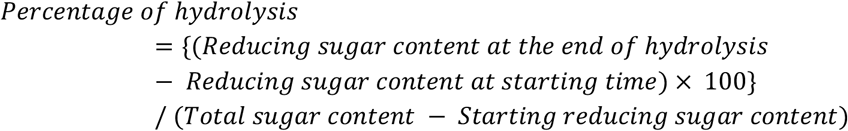

#### 2.7.2 Evaluation of prebiotic activity Score

A substance is classified as prebiotic when it promotes the proliferation of gut probiotics while suppressing the growth of pathogenic bacteria. The prebiotic potential of a molecule is assessed by determining whether the tested carbohydrate can sustain the growth of the probiotic as the exclusive sugar source. The prebiotic activity index of BMOs was calculated using the approach described by Kumar et al. (2020), with minor changes. Three strains of lactobacilli (*Lactobacillus acidophilus* MTCC 10307, *Lactobacillus rhamnosus* MTCC 1408, *Lactobacillus plantarum* MTCC 2621) were chosen as probiotic bacteria. In contrast, *Escherichia coli* ATCC 15223 was selected as a pathogenic bacterium. Lyophilised BMOs, FOS, and glucose were dissolved in incomplete MRS broth (without sugar source) for probiotics and M9 broth (lacking carbohydrates) for *E. coli* at a concentration of 1% (w/v). Incomplete MRS media and M9 media without carbohydrates were considered blank sets. 1% (v/v) (10^6 CFU/mL) of overnight specific probiotics and *E. coli* cultures were inoculated into each tube. The cultures were maintained at 37 °C. 10 μL of culture broth from each tube was collected and plated on agar plates. Culture colonies were counted on MRS agar plates (for probiotic strains) and nutrient agar plates (for *E. coli*) after 0 and 24 h of incubation. Prebiotic activity score was assessed utilizing the following formula:

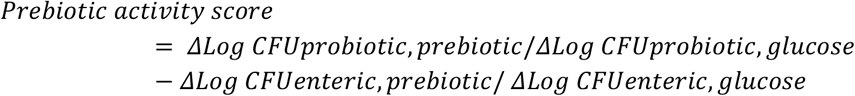

*Where, Δ Log CFU: the change in the logarithm of colony forming units(CFU) over time, specifically from* 0*th hour to* 24*th hour, probiotic, prebiotic*: *the grown of the probiotic bacteria on the prebiotic substrate, probiotic, glucose*: *the growth of the probiotic bacteria on the control substrate, gulcose, enteric,prebiotic* : *the growth of the enteric bacteria* (*E*.*coli*) *on the prebiotic substrate, enteric, glucose* : *the growth of the enteric bacteria on the control substrate, glucose*.

#### 2.7.3 Detection of SCFAs by HPLC

When probiotic bacteria are cultivated in the presence of prebiotics, short-chain fatty acids (SCFAs), specifically acetic, propionic, and butyric acids, are generated in the culture media. Three different probiotics (*Lactobacillus acidophilus* MTCC 10307, *Lactobacillus rhamnosus* MTCC 1408, *Lactobacillus plantarum* MTCC 2621) were grown in BMO-rich MRS medium at 37 °C for 48 h. After 24 and 48 h of incubation, 300 μL of media was taken out from each tube, centrifuged at 5000 g for 10 mins, and filtered through 0.45 μm filter membranes before injection (20 μL) into the HPLC (Agilent Infinity series 1200). A Hypersil Gold C18 column (250 × 4.6 mm, 5 μm) was used with a column temperature of 30 °C and a DAD detector set at 210 nm. As mobile phases, sodium dihydrogen phosphate (20 mM) buffer and acetonitrile were used. The HPLC separation programme was followed as described by Mitra et al. (2016). Individual SCFAs were quantified by interpolating the peak areas against standard calibration curves generated for each respective SCFA.

#### 2.7.4 Auto-aggregation assay of probiotics using BMOs

Auto-aggregation assay was carried out according to Yin et al. (2024) with some modifications. Each *Lactobacillus* strain was cultured for 24 h at 37 °C in different sugar-rich MRS medium (Glucose/FOS/BMOs + Incomplete MRS). The cells were harvested by centrifugation at 5000 g for 15 mins, washed twice, and resuspended in PBS to achieve approximately 10^6 CFU/mL. The cell suspensions were vortexed for 15 seconds, and auto-aggregation was evaluated at room temperature over 2 and 5 h. At 2 and 5 hours, 0.2 mL of suspension was collected from the upper layer of each tube and measured at 600 nm. The formula calculated the percentage of auto-aggregation of each probiotic:

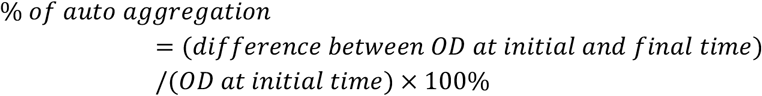

#### 2.7.5 Adhesion ability of probiotics to the HCT 116 cells

The adhesion of probiotics to HCT 116 cells in the presence of different carbohydrate standards and BMOs was assessed using the method reported by Megur et al. (2023) with some modifications. The actively growing HCT 116 cells were seeded into a sterile flat-bottom 12-well microtiter plate at a density of 10^5 cells per well and cultured at 37 °C with 5% CO_2_ until they reached 90% confluence. Before 24 hours of probiotic inoculation in the wells, the media were carefully replaced with antibiotic-free DMEM. After 24 hours, the spent media was gently aspirated to ensure the cells remained undisturbed. Glucose, FOS, and BMOs (1 mg/mL each) were separately mixed with serum-free, antibiotic-free DMEM, filtered with a 0.22 μm syringe filter, and added to respective wells. Actively growing cultures of *Lactobacillus plantarum, Lactobacillus acidophilus*, and *Lactobacillus rhamnosus* were mixed in a ratio of (1:1:1) and centrifuged at 5000 g for 5 mins. After centrifugation, the cell pellet was resuspended in 1 mL of serum-free, antibiotic-free DMEM media to achieve a final density of 10^6 CFU/mL. 100 μL of that probiotic mixture was added per well and incubated at 37 °C and 5% CO_2_ for 4 h. The well containing HCT 116 cells with bacteria was used as a control.

After incubation, the spent media were discarded, and the cell monolayer was rinsed three times with sterile PBS (pH 7.4) to eliminate non-adherent probiotics. To detach adherent cells from culture vessels, trypsin-EDTA solution was added and incubated for 10 mins. The suspension of adherent *lactobacillus* cells was serially diluted using saline solution and plated on an MRS agar plate. After 48 h of incubation at 37 °C, the number of colonies was counted, and the percentage of adhesion was calculated by the following equation:

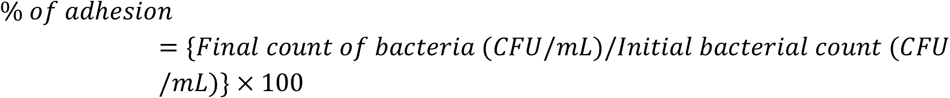

### 2.8 Statistical analysis

All of the experimental data was analysed using Microsoft Excel Office 2019. All experiments were performed in three copies, and the data are presented as mean ± standard deviation. The statistical significance (p value) was calculated by performing two-sample, unpaired Student’s t-tests (*p < 0.05, **p < 0.01, ***p < 0.001).

## 3. Results and discussion

### 3.1 Isolation and partial purification of BMOs

Upon adjusting the extraction parameters (solvent for isolation, solvent concentration, sample-to-solvent ratio, temperature, and extraction time) using a one-factor-at-a-time approach, the maximum low-molecular-weight carbohydrates were extracted from barnyard millet flour (particle size between 500 and 150 μm) with 80% (v/v) aqueous ethanol at 60°C for 1 h at a 1:5 sample-to-solvent ratio. Many conventional methods for isolating lower-chain-length carbohydrates use a higher alcohol concentration (Ekvall et al., 2006). With those optimized parameters, the yield of the crude extracted sugar from barnyard millet flour was 3.61±0.24% of the total carbohydrate content. The yield was lower than that of the crude oligosaccharides extracted from pearl millet (Mondal et al., 2022). It has been reported that wheat, oat, barley, and rye contain 0.478%, 0.083%, 0.236%, and 0.081% of raffinose family oligosaccharides (RFOs), respectively (Henry & Saini, 1989), and Shewry and Hey (2015) noted that the fructan content in wheat can vary from 0.84% to 2.3%. So, the total crude low molecular weight carbohydrates (DP<10) content in barnyard millet grain is higher than the percentage of total RFOs and fructan content in whole wheat grain.

The crude extracted oligosaccharides were further purified through adsorption column chromatography. The TLC profile (Fig. 1I) of charcoal-column eluted sugar shows that monosaccharides (R_f_ ≥ R_f_ value of glucose) were maximally eluted with 5% aqueous ethanol (v/v). With the increasing aqueous ethanol percentage (%) (v/v), sugars with longer chain length started to elute. From a 15% ethanol wash, fewer disaccharides (R_f_ > R_f_ value of raffinose) were eluted. Most oligosaccharides were eluted with multiple 15% and 20% ethanol washes. From the elution profile (Fig. 1III) using 15% to 30% aqueous ethanol as a mobile phase on a charcoal-celite column, the recovery of oligosaccharides was 56.41±1.67% of crude extract with 85% of purity. These findings align with the data presented by many authors (Kuhn & Filho, 2010; Mondal et al., 2022). Although there was only a small amount of protein in the crude extract, 89.54±0.91% of the protein was removed after it eluted through the adsorption column chromatography. It has been determined that the barnyard millet extract lacked phenolics following elution from the activated charcoal column (Fig. 1IV,1V). Commercial activated carbon successfully removed phenolic compounds from liquid-soaked distilled grape pomace, as shown by Soto et al. (2007). Campos et al. (2017) reported that FOS was free from the predominant residual turbidity and pigments after treatment with activated carbon. A purity of 94.23% was achieved for the extracted oligosaccharides after dialysis. Fig. 1VI indicates that almost all disaccharides were removed after dialysis.

**Fig. 1:**
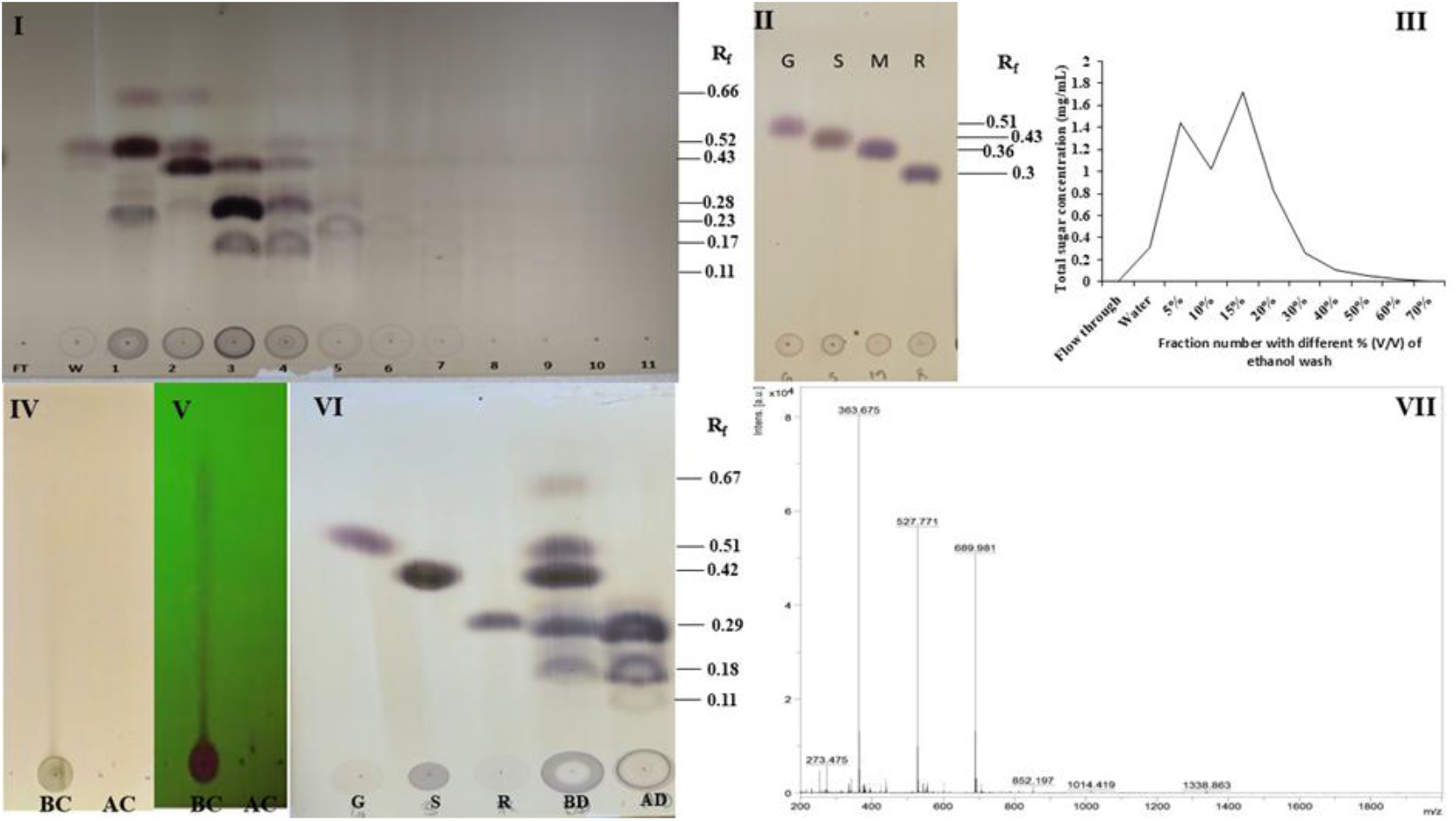
TLC profile of charcoal-column eluted samples, FT-flow through, W-water, 1-5% aqueous (aq.) ethanol (v/v) run, 2-10% aq. ethanol (v/v) run, 3-15% aq. ethanol (v/v) run, 4-20% aq. ethanol (v/v) run, 5-30% aq. ethanol (v/v) run, 6-40% aq. ethanol (v/v) run, 7-50% aq. ethanol (v/v), 8-60% aq. ethanol (v/v) run, 9-70% aq. ethanol (v/v) run, 10-80% aq. ethanol (v/v) run, 11-90% aq. ethanol (v/v) run, **II**: TLC profile of charcoal-column eluted standards, G-glucose, S-sucrose, M-maltose, R-raffinose, **III**: Adsorption chromatogram showing sugar elution profile with increasing percentage of aqueous ethanol, **IV**: TLC profile of phenolic compounds in extracted oligosaccharides staining with 2% ferric chloride (ethanol) and **V**: under UV (254nm), BC-before charcoal adsorption chromatography, AC-after charcoal adsorption chromatography, **VI**: TLC profile of oligosaccharides before and after dialysis, G-glucose S-sucrose, R-raffinose, BD-before dialysis, AD-after dialysis, **VII**-MALDI-TOF profile of the extracted oligosaccharides.

### 3.2 Characterization of BMOs by MALDI-TOF

Several significant peaks at m/z values of 363, 527, 689, 852, 1014, and 1338 were detected in the MALDI-TOF (Fig. 1VII). The sodium ion (Na^+^) helped with the detection in the time-of-flight (TOF) experiment, and each mass-to-charge (m/z) value corresponds to a charged oligosaccharide ion ([oligosaccharide + Na]^+^). The analysis of the m/z progression revealed that the obtained oligosaccharides had degrees of polymerization ranging from 2 to 8. Mondal et al. (2022) reported that extracted pearl millet oligosaccharides have the same chain length range, and the calculated mass difference between consecutive peaks reflects the molecular weight of a single hexose monosaccharide unit. Furthermore, the molecular weights are comparable to those of β-Glucooligosaccharides found in oats, as stated by Jiang et al. (2025).

### 3.3 Thermal properties of BMOs

The thermal degradation behavior of the oligosaccharide mixture was studied across a temperature range of 35 to 600 °C. The TGA thermogram (Fig. 2) showed three different areas of mass loss with increasing temperature: 35 to 150 °C, 150 to 400 °C, and 400 to 600 °C. The weight reduction in the first zone (35–150 °C) was attributed to the evaporation of water as the temperature increased. The method enables the quantification of the water content in the samples by calculating the weight loss from the TGA signal, which was nearly 10%. A similar result was also reported in pectic oligosaccharides (Combo et al., 2012). The second thermal degradation process, from 150 to 400 °C, was characterised by a quick mass loss. This substantial decrease was probably related to the breakdown of the oligosaccharide backbone through the cleavage of glycosidic linkages. Völkel et al. (2022) noted that one of the causes of unexpected crosslinking from heat affecting polymeric cellulose was thermal breakdown of glycosidic linkages. The third area shows a slow, steady mass loss between 400 and 600 °C. Within this temperature range, the polymeric molecules break down due to heat, resulting in the formation of volatile gases. The TGA DTA profile of BMOs is very close to the thermal profile of mannooligosaccharides reported by Magengelele et al. (2023).

**Fig. 2:**
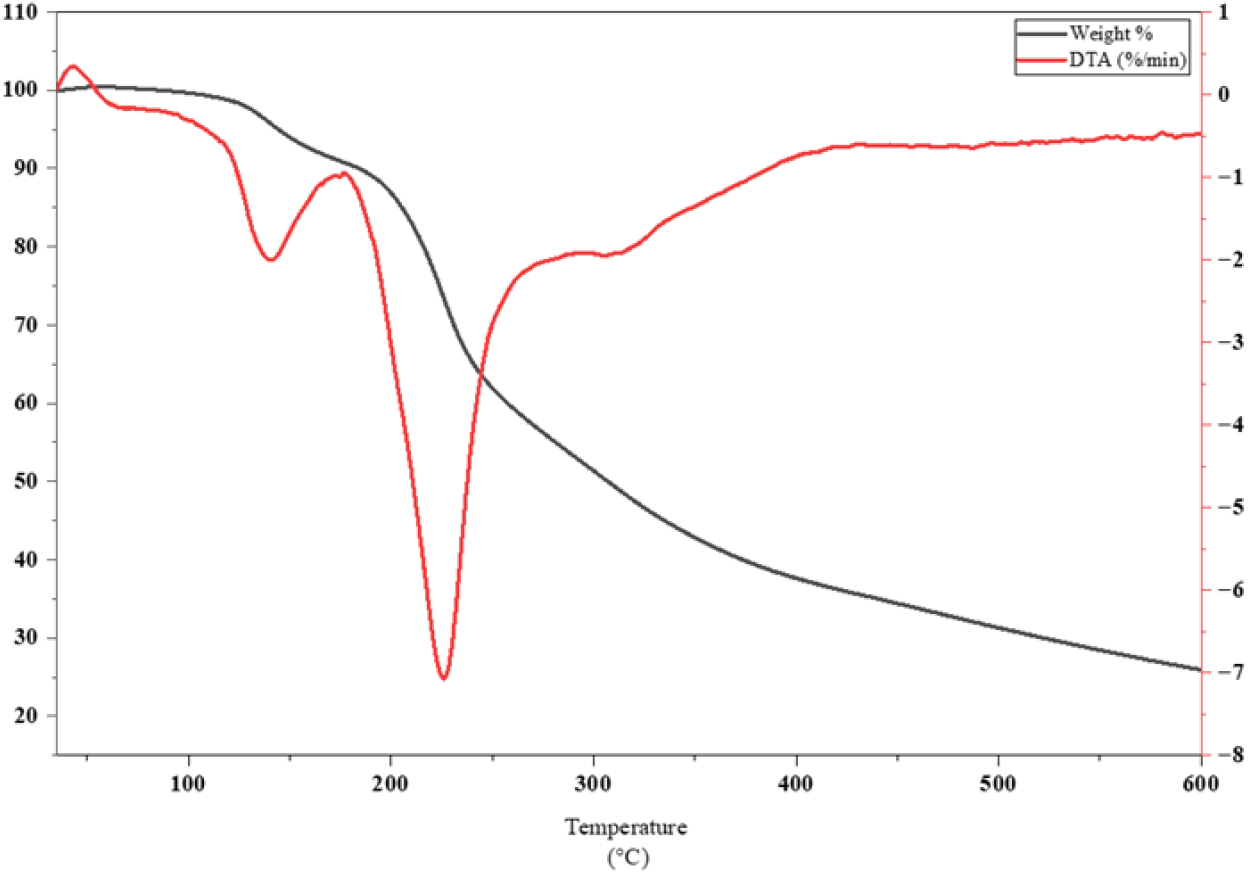
TGA and DTA curve of extracted barnyard millet oligosaccharides.

### 3.4 Prebiotic potential of BMOs

#### 3.4.1 Non-digestibility Assay of BMOs

Any molecule is considered prebiotic if it is non-digestible by mammalian α enzymes, stomach acid, and goes directly to the large intestine. In the *in vitro* system, the non-digestibility of barnyard millet oligosaccharides was investigated by treating the molecules with salivary α-amylase, porcine pancreatic α-amylase, and artificial gastric juice. After 6 h of treatment, it was found that in each case, the digestibility percentage of BMOs was significantly lower (the percentage of hydrolysis of BMOs for salivary α-amylase, pancreatic α-amylase, and artificial gastric juice was 2.3±0.18, 6.45±0.49, and 0.89±0.27%, respectively) compared to starch and very close to FOS (Fig. 3). Therefore, we can say that approximately 91% of BMOs are not digested by intestinal enzymes and will pass directly to the colon. Dasaesamoh et al. (2016) also reported that 87.59% of dragon fruit oligosaccharides reach the large intestine after passing through the upper gastrointestinal tract. Various studies indicate that galactooligosaccharides (GOS) are approximately 90% undigested upon reaching the colon (Ambrogi et al., 2021). These reported observations are consistent with the findings of the present study. Salivary α-amylase and pancreatic α-amylase can hydrolyse (α-1,4) glycosidic bonds of starch and break them into small monosaccharides like glucose or oligosaccharides like maltose. So, we can predict that the internal structure of BMOs may consist of branched or β-linked glycosidic bonds.

**Fig. 3:**
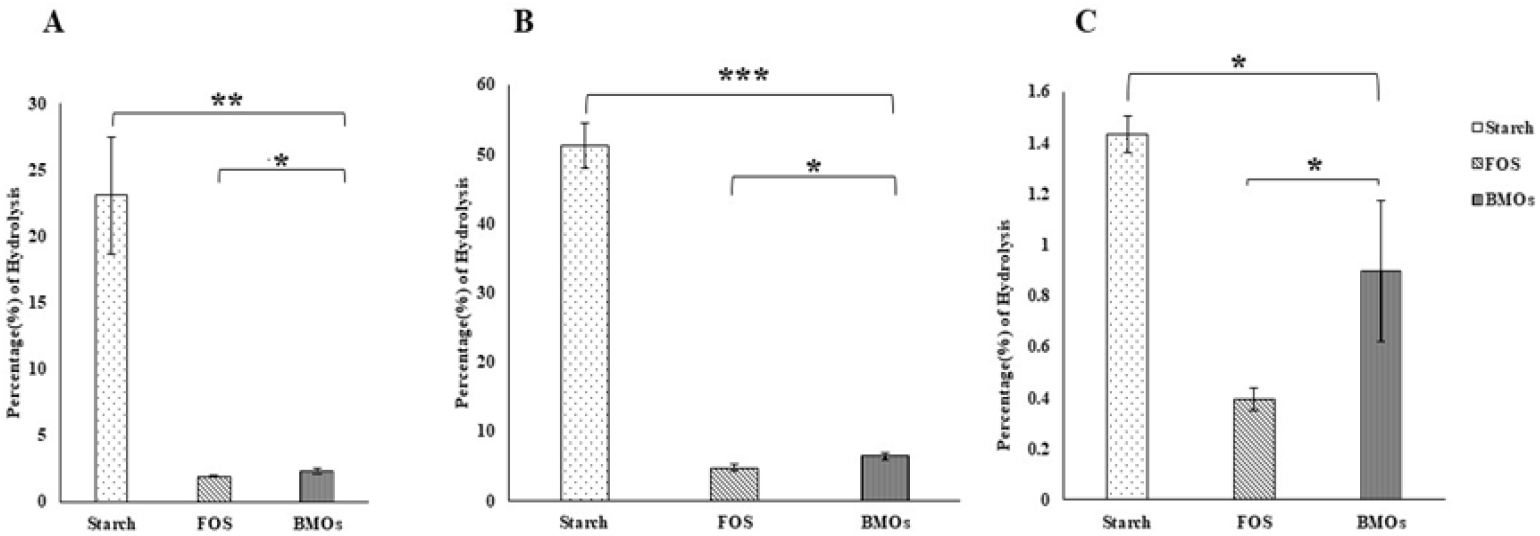
*In vitro* non-digestibility assay of BMOs, hydrolysis percentage achieved using **A**-human salivary alpha(α)-amylase, **B**-porcine pancreatic alpha (α)-amylase, and **C**-gastric pH. Data are presented as the mean ± standard deviations (n=3). The statistical significance (p value) was calculated by performing two-sample, unpaired Student’s t-tests (*p < 0.05, **p < 0.01, ***p < 0.001).

#### 4.3.2 Assessment of prebiotic activity score

The prebiotic activity score, using BMOs as the sole carbohydrate source, was assessed by measuring the growth of three distinct lactobacilli strains compared with pathogenic bacteria. From the prebiotic activity score formula, a positive score indicates that the growth of probiotics on the examined prebiotic as the exclusive carbohydrate source is higher than the growth of the pathogen (*E. coli*) on the same prebiotic-rich media compared with glucose-rich media. Considering the established metabolic diversity of lactobacilli, significant variance in scores for individual prebiotic molecules was anticipated when used by other probiotic strains. The prebiotic activity scores of BMOs compared with other prebiotic FOS with respect to three different lactobacilli (*L. acidophilus, L. plantarum, L. rhamnosus*) are shown in Figure 4. The highest score was seen in *Lactobacillus rhamnosus*, followed by *Lactobacillus plantarum*, with values of 0.88 ± 0.13 and 0.66 ± 0.04, respectively. *Lactobacillus acidophilus* preferentially digested FOS over BMOs, as indicated by the prebiotic activity score of 0.48 ± 0.03 for FOS, in contrast to 0.36 ± 0.04 for BMOs. Huebner et al. (2007) reported that GOS has a prebiotic activity index of 0.82 on *L. plantarum* 4008. Similarly, Kumar et al. (2020) observed that laminari-oligosaccharides (LOS) significantly enhanced the growth of probiotics, yielding prebiotic activity scores of 0.92 ± 0.01 for *L. plantarum* DM5. Therefore, the strain’s specific activity scores determine the ability to utilize β-linked or complex oligosaccharides. Considering the non-digestibility assay of the BMOs, these positive prebiotic activity scores indicate that BMOs exhibit the basic characteristics of a selected prebiotic carbohydrate.

**Fig. 4:**
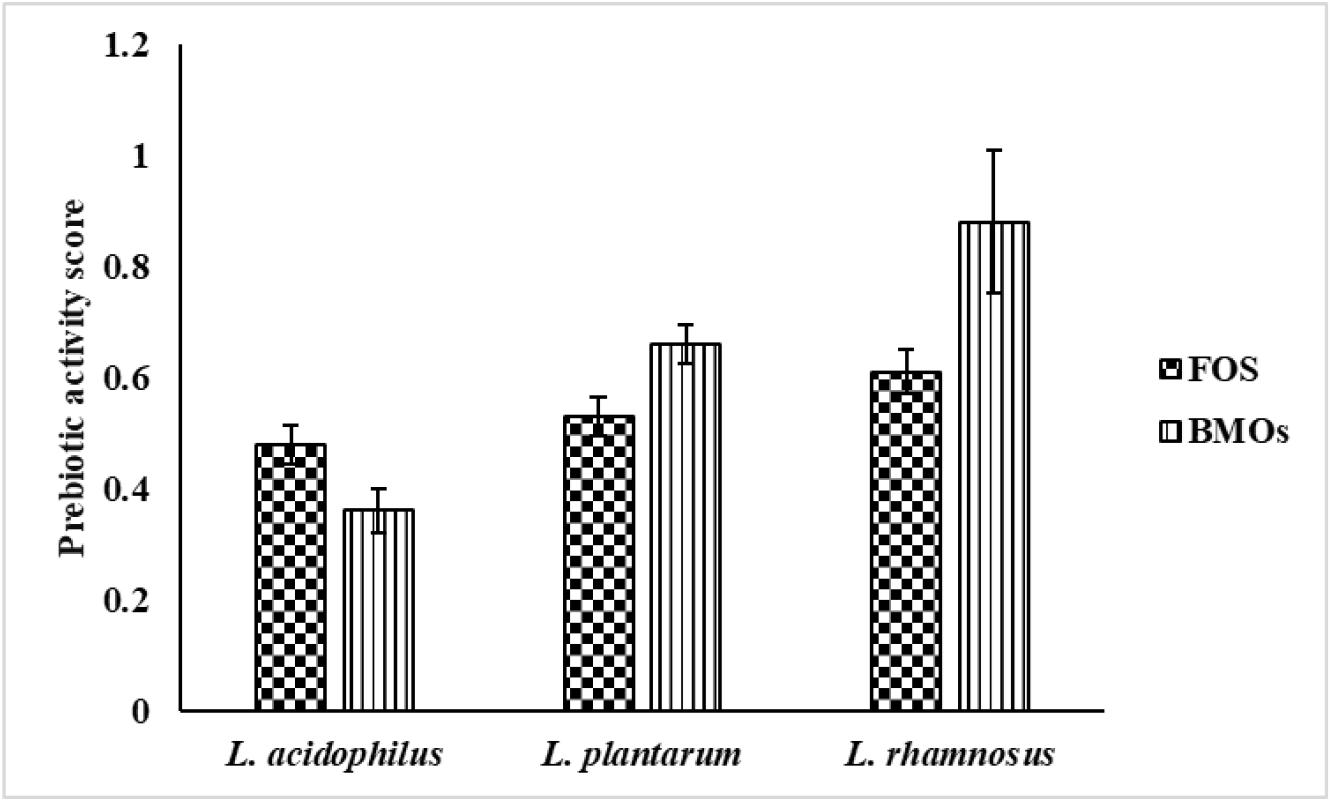
Graphical depiction of prebiotic activity scores across various probiotic bacteria on FOS and BMOs. Values are expressed as the mean ± standard deviations (n=3).

#### 4.3.3 Detection of SCFAs by HPLC

By using the prebiotic as the sole carbohydrate source, probiotics, especially *lactobacillus sp*., can produce SCFAs, which inhibit the growth of pathogenic microbes by lowering gut pH (Yan et al., 2024) and reinforcing the intestinal barrier, enhancing nutritional absorption (Ca, Fe, Mg) (Markowiak-Kopeć & Śliżewska, 2020). Tan et al. (2014) reported that receptors (especially GPCRs) for short-chain fatty acids play a significant role in the control of metabolic and inflammatory responses. SCFAs produced by the *in vitro* digestibility of BMOs by different probiotic Lactobacilli were assessed using HPLC. The amounts of SCFAs and lactic acid produced by different strains using BMOs after 24 and 48 h are presented in Table 1. *L. rhamnosus* produced the highest amount of acetic acid, butyric acid, formic acid, and propionic acid with 6.82±3.23, 4.37±0.28, 3.37±0.87, and 1.22±0.27 mg/mL (p ≤ 0.05), respectively, after 48 h of fermentation. The maximum amount of propionic acid and formic acid (0.94±0.08, 2.24±0.17 mg/mL) was produced by *L. acidophilus* after 24 h. At the same time (at 24 h of fermentation), *L. plantarum* produced almost the same amount of formic acid as *L. acidophilus*. The amount of lactic acid was found to be the highest in *L. plantarum* (2.54±2.7 mg/mL) after 48 h, compared with other strains. However, *L. plantarum* produced no butyric acid up to 48 h of fermentation using BMOs as prebiotics.

**Table 1:** SCFAs produced by various probiotic strains using BMOs as carbohydrate source. Values are expressed as the mean ± standard deviations (n=3). Differences in values for a specific parameter are considered statistically significant. (Two-way ANOVA t-test, p ≤ 0.05)

| 24 h |  |  |  |
| --- | --- | --- | --- |
| SCFA<br>(mg/mL) | *L. acidophilus* | *L. plantarum* | *L. rhamnosus* |
| Acetic acid | 1.32±0.16 | 2±1 | 3.35±2.53 |
| Butyric acid | 1±0.24 | 0 | 2.98±2.57 |
| Propionic acid | 0.94±0.08 | 0.16±0.05 | 0.74±0.25 |
| Formic acid | 2.24±0.17 | 2.24±0.7 | 1.59±0.07 |
| Lactic acid | 0.17±0.01 | 0.92±0.42 | 0.99±0.09 |
| 48 h |  |  |  |
| SCFA<br>(mg/mL) | *L. acidophilus* | *L. plantarum* | *L. rhamnosus* |
| Acetic acid | 1.63±0.37 | 3.03±2.9 | 6.82±3.23 |
| Butyric acid | 1.41±0.32 | 0 | 4.37±0.28 |
| Propionic acid | 0.99±0.1 | 0.3±0.15 | 1.22±0.27 |
| Formic acid | 2.6±0.31 | 2.7±1.51 | 3.37±0.87 |
| Lactic acid | 0.62±0.6 | 2.54±2.7 | 1.38±0.18 |

The study on lactic acid and short-chain fatty acids (SCFAs) demonstrated that the specific type and quantity of SCFAs produced by a particular probiotic were reliant upon the kind of carbohydrates fermented. A notable enhancement of lactic acid, acetic acid, and propionic acid after fermentation by both *L. rhamnosus* and *L. acidophilus* with GOS (DP 3-5) as the only carbon source of the growth medium was reported by Shi et al. (2018). In our experiment, the prebiotic activity score of *L. acidophilus* was lower than that of *L. plantarum*, but *L. acidophilus* produced more butyric and propionic acids after fermentation when using the BMOs as a carbohydrate source. These strain-specific, substrate-dependent patterns may be due to the utilization of carbon sources via different pathways (e.g., propionogenic, butyrogenic, etc.). The divergence between the prebiotic index and SCFA yield may depend on the strains’ biomass and metabolite outputs (Agrawal et al., 2026). The SCFA pattern observed in this work thus supports the possible existence of BMOs as highly active metabolic prebiotic substrates that can influence intestinal fermentation.

#### 4.3.4 Auto aggregation assay

Auto-aggregation causes biofilm to form in the gastrointestinal system. This protects the intestine by creating a barrier and boosting immunity, while also preventing the growth of harmful bacteria (B et al., 2019). In our study, we performed an auto-aggregation assay using three probiotics (*L. acidophilus, L. plantarum*, and *L. rhamnosus*) on BMOs, comparing them with glucose and FOS. As depicted in Fig. 5, the auto-aggregation percentages of three different lactobacilli were increased in the presence of FOS (% of auto-aggregation were 20.56±0.53, 24.67±1.37, 29.74±2.52% (p ≤ 0.05) in case of *L. acidophilus, L. plantarum*, and *L. rhamnosus*, respectively) and BMOs (% of auto-aggregation were 20.58±0.42, 26.64±2.12, 30.78±3.02% (p ≤ 0.05) in case of *L. acidophilus, L. plantarum*, and *L. rhamnosus*, respectively) compared with glucose after 5 h. The highest percentage of autoaggregation occurred after 5 h using BMOs in *L. rhamnosus*, with an increase of 6.44% compared to glucose. Increased auto-aggregation of probiotic bacteria leads to greater colonization of the gut wall, thereby preventing adhesion of pathogenic bacteria (Tareb et al., 2013). Similarly, Yin et al. (2024) reported that isomalto-oligosaccharides (IMO) and FOS significantly promoted the auto-aggregation rates of *L. plantarum* and *L. rhamnosus* strains. A significant aspect affecting a probiotic strain’s adhesion to the gastrointestinal tract is its capacity for auto-aggregation (Kiran et al., 2024). In our study, an increasing percentage of auto-aggregation of probiotics using BMOs suggests that this prebiotic acts as an environmental trigger that encourages bacteria to clump together, which is a prerequisite for effective colonization and competitive exclusion of pathogens.

**Fig. 5:**
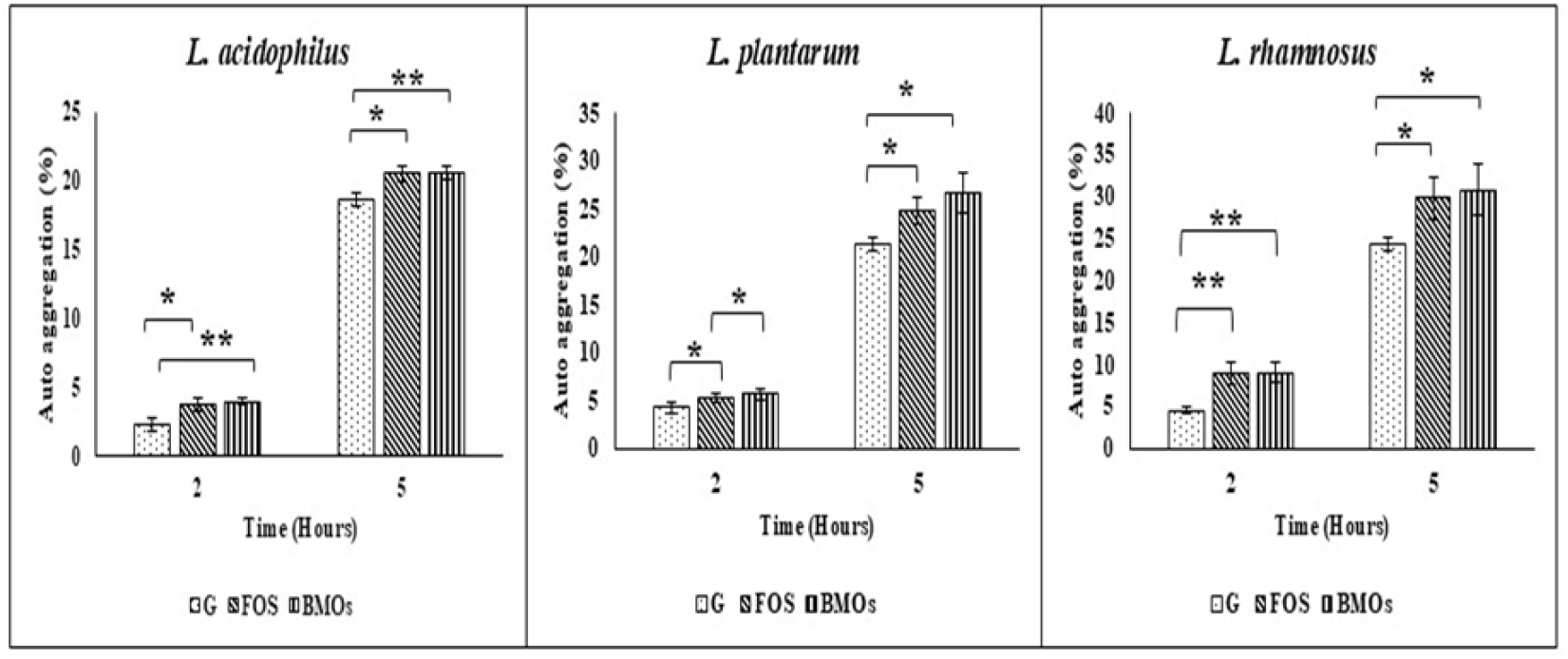
Auto-aggregation percentages of different Lactobacilli in the presence of different carbohydrates, including BMOs. Values are shown as the mean ± standard deviations (n=3). The statistical significance (p value) was calculated by performing two-sample, unpaired Student’s t-tests (*p < 0.05, **p < 0.01).

#### 4.3.5 Adhesion ability of probiotics to the HCT 116 cells

The HCT 116 cell line exhibits structural and functional similarities to mature intestinal epithelial cells (Megur et al., 2023). Consequently, we used this cellular model to determine the adhesion properties of different Lactobacilli in the presence of FOS, glucose, and BMOs. The highest adhesion percentage (18.06±1.16%) of *lactobacillus* strains to HCT 116 cells occurred in the presence of BMOs (Fig. 6). The FOS and glucose showed adhesion percentages of 14.83±1.7% and 3.54±1.48%, respectively, compared with the control (Cell + Bacteria). *Lactobacillus* cells adhered to HCT 116 cells more efficiently (5.1-fold) in the presence of BMOs than in the presence of glucose. Moreover, increased adherence of probiotic strains to HCT 116 cells upon treatment with BMOs indicates their support for probiotic colonization in the gut *in vivo* as well.

**Fig. 6:**
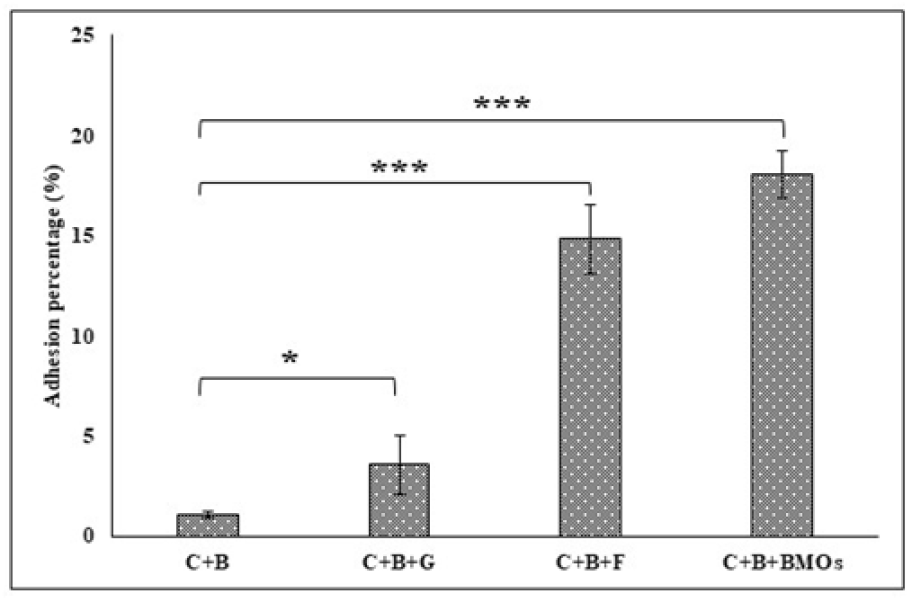
Adhesion percentages of different probiotics to HCT116 cells in the presence of various carbohydrates. (C-HCT 116 cells, B-Mixer of probiotic bacteria, F-FOS, G-Glucose, BMOs-Barnyard millet oligosaccharides). Data are represented as the mean ± standard deviations (n=3). All the data are significantly different (p ≤ 0.05).

## 5. Conclusion

Although various health benefits of taking barnyard millet were reported in different literature, till now, the content of non-digestible oligosaccharides present in it and its prebiotic activity have not been explored. In our study, crude low-molecular-weight carbohydrates (including oligosaccharides) were extracted from barnyard millet under optimal extraction conditions (80% aqueous ethanol (v/v) with a 1:5 sample, solvent ratio for 1 h at 60 °C), yielding 3.61±0.24%. All monosaccharides, maximum disaccharides, phenolics, and most of the proteins were removed from the oligosaccharides by passing the crude extracts through adsorption chromatography (charcoal-celite column). The MALDI-TOF mass spectrum showed that the partially purified crude extract contained different oligosaccharides (degrees of polymerization ranging from 2 to 8). BMOs were free of other low-molecular-weight carbohydrates by dialysis through dialysis tubes (MWCO 500D). Non-digestibility assays indicated that approximately 91% of BMOs were not digested by intestinal enzymes and passed directly to the lower intestine. From the prebiotic activity score, the strain-specific growth was observed, and *L. rhamnosus* showed the highest prebiotic score (0.88±0.13) utilizing BMOs as the sole carbohydrate source. Also, these oligosaccharides helped *Lactobacillus sp*. make lactic acid and SCFAs. In addition, BMOs increased the auto-aggregation percentages of *Lactobacillus* strains. In the presence of BMOs, Lactobacilli were observed to be more adherent to HCT116 cells. Hence, these results suggest that BMO could be a good prebiotic candidate. More research is needed in artificial gut models or *in vivo* studies to fully understand its prebiotic efficacy and anti-diabetic potential, as well as to support its use in functional foods and nutraceutical products that promote gut health.

## Credit authorship contribution statement

**Sachin Maji**: Conceptualization, Methodology, Investigation, Formal analysis, Writing – Original Draft. **Paramita Biswas**: Methodology. **Shivangi Agrawal**: Writing – review & editing. **Sandip Shit**: Writing – review & editing. **Satyahari Dey**: Project administration, Supervision, Writing – review & editing.

## Acknowledgements

The financial assistance from the Ministry of Education, Government of India, is sincerely appreciated. We endorse the Central Research Facility (CRF) at IIT Kharagpur for supplying essential infrastructural and analytical support.

## Declaration of interests

The authors declare that they have no known competing financial interests or personal relationships that could have appeared to influence the work reported in this paper.

## Abbreviation

v/v: volume/volume
FOS: Fructooligosaccharides
XOS: Xylooligosaccharides
GI: Gastrointestinal
MWCO: Molecular Weight Cut-Off
w/v: weight/volume
MALDI-TOF: Matrix-Assisted Laser Desorption Ionization—Time of Flight
MRS: De Man-Rogosa-Sharpe
HPLC: High-Performance Liquid Chromatography
DAD: Diode Array Detector
DMEM: Dulbecco’s Modified Eagle Medium

## Notes

### Competing Interest Statement

The authors have declared no competing interest.

